# Role of PML-Nuclear Bodies in Human Herpesvirus 6A and 6B Genome Integration

**DOI:** 10.1101/413575

**Authors:** Vanessa Collin, Annie Gravel, Benedikt B. Kaufer, Louis Flamand

## Abstract

Human herpesviruses 6A and 6B (HHV-6A/B) are two betaherpesviruses that readily integrate their genomes into the telomeres of human chromosomes. To date, the cellular or viral proteins that facilitate HHV-6A/B integration remain elusive. In the present study, we demonstrate that the immediate early protein 1 (IE1) of HHV-6A/B colocalizes with telomeres during infection. Moreover, IE1 associates with PML-NBs, a nuclear complex that regulates multiples cellular mechanism including DNA repair and antiviral responses. Furthermore, we could demonstrate that IE1 targets all PML isoforms and that both proteins colocalize at telomeres. To determine the role of PML in HHV-6A/B integration, we generated PML knockout cell lines using CRISPR/Cas9. Intriguingly, in the absence of PML, the IE1 protein could still localize to telomeres albeit less frequently. More importantly, HHV-6A/B integration was impaired in the absence of PML, indicating that it plays a role in the integration process. Taken together, we identified the first cellular protein that aids in the integration of HHV-6A/B and shed light on this targeted integration mechanism.

**Author summary:** Human herpesviruses type 6A and 6B are relatively common viruses whose infections can be life threatening in patients with a compromised immune system. A rather unique feature of these viruses is their ability to integrate their genome in human chromosomes. Integration takes place is a specialized region of the chromosomes known as telomeres, a region that controls cellular lifespan. To date, the mechanisms leading to HHV-6A and HHV-6B integration remain elusive. Our laboratory has identified that the IE1 protein of HHV-6A and HHV-6B target the telomeres. Moreover, we have shown that IE1 associates with a cellular protein, PML, that is responsible for the regulation of important cellular mechanisms such as the life span of cells and DNA repair. Hence, we studied the role of PML in HHV-6 integration. Our study demonstrates that in absence of PML, the HHV-6A and HHV-6B integrate 50-70% less frequently. Thus, our study unveils the first cellular protein involved in HHV-6A and HHV-6 chromosomal integration.

## Introduction

Human herpesviruses type 6A and 6B (HHV-6A/B) are members of the *betaherpesvirinae* that were isolated in the 1980’s. In 2013, the International Committee on Taxonomy of Viruses recognized HHV-6A and HHV-6B as distinct viral species (1). HHV-6B is known as the etiologic agent of *exanthem subitum,* a childhood disease whose symptoms include, fever, occasional skin rash and respiratory distress (2). HHV-6A is much less characterized than HHV-6B. Considering that many HHV-6A/B proteins share 90-95% homology, the symptoms of primary HHV-6A infections are likely lessened in part due to cross-protective immunity developed against HHV-6B. Upon primary infection, HHV-6A/B establish latency like all herpesviruses. During latency, most herpesviruses maintain their genome in a circularized form (episome). The viral episomes are generally tethered to the human chromosomes, ensuring transmission to daughter cells following cell division (3), (4). However, to date, the presence of HHV-6A/B episomes during latency have yet to been demonstrated.

Despite the fact that no episomes of HHV-6A/B have been reported, both viruses can reactivate and cause secondary infections. In healthy subjects, HHV-6B reactivation is mostly subclinical and controlled by the immune system. However, in immunocompromised individuals, HHV-6B reactivation can be problematic and occasionally life-threatening (5),(6),(7). In case of HHV-6A, several reports have associated the virus with neurodegenerative diseases such as multiple sclerosis and more recently with Alzheimer’s disease (8), (9), (10), (11). In spite of their pathological differences, both HHV-6A and HHV6B can readily integrate their genomes into host chromosomes (12), (13), (14). HHV-6A/B integration can take place in various chromosomes but invariably occurs within the telomeric region (12), (13), (14). In 1993, Luppi et *al* reported three cases of individuals with telomeric integration of HHV-6A/B (13). In 1999, Daibata et *al* subsequently demonstrated that chromosomally-integrated HHV-6 can be inherited (12). Subjects with inherited chromosomally-integrated HHV-6A/B (iciHHV-6A/B) have at least one (occasionally 2 or 3) copy of the viral genome present in every somatic cells, with the viral genome transmitted to 50% to their children (15), (16). Viral integration into telomeres could be an alternative latency mechanism for HHV-6A/B. In support, the integrated HHV-6A/B genomes are generally intact and conserved without any gross rearrangements or mutations (17). Furthermore, integrated HHV-6A/B genomes can express genes and lead to complete viral reactivation (18)(19),(20),(7). Reactivation of HHV-6A/B can be life threatening for immunocompromised hosts.

Telomeres are non-coding (TTAGGG)_n_ hexanucleotides present at the chromosome termini and contain a single-stranded 3’ extension of 30-500 G rich nucleotides. They protect chromosomes against the loss of genetic information, which would result in premature cell senescence and prevent the recognition of chromosome ends by the DNA damage response (DDR) machinery. The telomere end forms a t-loop (22),(23) that is maintained by a complex of 6 proteins, the shelterin proteins (24), (25) that protect the chromosomes against DNA damage response. The HHV-6A/B genome is about 160 kilobase pairs (kbp) in length and contains a unique region (U) with close to 100 open reading frames (26), (27), (28). This U region is flanked by identical direct repeat regions (DR_L_ and DR_R_) of 8-9 kbp that contain telomere sequences identical to human telomeres at both ends (27), (29). Wallaschek and al. recently demonstrated that these telomeric sequences facilitate integration of HHV-6A into host telomeres (30). This indicated that integration is likely mediated by homologous DNA recombination events. To date, no viral or cellular proteins have been identified that are involved in HHV-6A/B integration.

An interesting candidate involved in viral integration is the immediate-early protein 1 (IE1) of HHV-6A/B, which can be transcribed without *de novo* protein synthesis (31). The IE proteins of herpesviruses regulate early genes and plays an important role in the initiation of lytic virus replication. Moreover, they establish a favorable environment by manipulating PML-Nuclear bodies (PML-NBs), which are part of the cellular antiviral defense (32), (33), (34). In the context of an infection, PML-NBs have been shown to repress replication of various viruses with its components SP-100 and DAXX. PML-NBs are found mostly in the nucleus and contain large quantities of the PML protein (35), (36). Some viruses have developed ways to overcome this antiviral mechanism by degrading or manipulating PML-NBs. For example, herpes simplex virus 1 (HSV-1) encodes the E3 ligase ICP0 that conjugates ubiquitin to PML and induces its degradation (37), (38). Human cytomegalovirus (hCMV) IE1 de-SUMOylates PML-NBs resulting in PML redistribution (39). In contrast, HHV-6A/B infection does induce dispersal of PML-NBs but reduces and increases their size (32), (33), (40). Intriguingly, HHV-6B IE1 has been shown to colocalize with PML during infection (32), (33) however, the role of this PML-IE1 interaction remains unknown.

Considering that 1) PML is located at telomeres, 2) PML-NBs associate with proteins involved in homologous recombination and 3) viral integration occurs at telomeres, we hypothesize that PML likely plays a role in HHV-6A/B chromosomal integration. We addressed this hypothesis and could demonstrate that HHV-6A/B IE1 not only localizes with PML, but also the host telomeres. In addition, we could demonstrate that PML indeed plays a role in HHV-6A/B integration.

## Materials and methods

### Cell lines and viruses

HeLa cells with long telomeres (Hela LT) (41) and HEK293T (ATCC, Manassas, VA, USA) were cultured in Dulbecco’s modified Eagle’s medium (DMEM; Corning Cellgro, Manassas, VA, USA) supplemented with 10% fetal bovine serum (FBS) (Thermo Fisher Scientific, Waltham, MA, USA), nonessential amino acids (Corning Cellgro), HEPES, sodium pyruvate (Wisent Inc., St-Bruno, Québec, Canada), and 5 μg/ml plasmocin (Invitrogen, San Diego, CA, USA). U2OS (osteosarcoma) cells (ATCC) were cultured in the same medium but supplement with 10% of Nu serum (Corning Cellgro) instead of FBS.

### Plasmids

Expression vectors for HHV-6A IE1 (pCDNA4/TO-IE1A) and HHV-6B IE1 (pCDNA4/TO-IE1B) control vector (pCDNA4/TO) were described previously (42). Plasmids expressing PML isoforms were kindly provided by Jin-Hyun Ahn (43). To generate a PML-I lentiviral vector, the PML-1 gene was PCR amplified with *attB1* and *attB2* sites added to the forward and reverse primer, respectively. The PCR amplicon was recombined into pDonor221 vector followed by a second recombination into pLenti CMV Hygro DEST vector, a kind gift from Eric Campeau and Paul Kaufman (Addgene plasmid # 17454) (44). The PML Double Nickase Plasmids (h2) (sc-400145-NIC-2) were bought from Santa Cruz Biotechnology (Santa Cruz, CA, USA).

### Immunofluorescence (IFA)

Coverslips were incubated for 30 minutes in blocking solution (1 mg/ml BSA; 3% goat serum; 0.1% Triton X-100; 1 mM EDTA pH 8.0, in phosphate-buffered saline (PBS)). After blocking, coverslips were incubated for 1 hour in primary antibody diluted in blocking solution. Coverslips were washed with PBS, three times for five minutes. Coverslips were incubated for 30 minutes with secondary antibody diluted in blocking solution. Coverslips were washed with PBS, three times for five minutes. When the IFA was done, coverslips were air dried at room temperature and mounted with *SlowFade* Gold Antifade reagent containing DAPI (Invitrogen, Eugene, Oregon USA).

### Immunofluorescence conjugated to *in situ* hybridization (IF-FISH)

Cells on coverslips were stained as for IFA. Once IFA was completed, cells were fixed for 2 minutes at room temperature with 1% paraformaldehyde in PBS. Coverslips were washed two times for five minutes with PBS. Cells were dehydrated with 5 minutes each consecutive ethanol baths (70%, 95%, 100%). Once dried, coverslips were placed upside down on a drop of hybridizing solution (70% formamide; 0.5% blocking reagent; 10 mM Tris-HCl pH 7.2; 1/1000 Cy3 or Cy5-TelC PNA probe). Samples were denatured for 10 minutes at 80°C on a heated block. Coverslips were incubated over night at 4°C in the dark and washed two times for 15 minutes in washing solution (70% formamide; 10 mM Tris-HCl pH 7.2). Coverslips were washed 3 time for 5 minutes with PBS and were air dried, slow fade was added and coverslips were sealed.

### Transfection assays

U2OS cells were seeded at 2 × 10^5^ cells/well in a 6-well plate containing glass coverslips in 2 mL of medium. Cells were transfected 24 hours post-seeding with pCDNA4/TO, pCDNA4/TO-IE1A, pCDNA4/TO-IE1B expression vector using the *TransIT*-LT1 Transfection Reagent (Mirus Bio LLC, Madison, WI, USA). After 48 hours of transfection, cells were washed 3 times with PBS and fixed in 2% of paraformaldehyde and used for immunofluorescence (IFA) assay. HeLa LT cells were seeded at 1 × 10^5^ cells/well in a 6-well plate containing glass coverslips in 2 mL of medium. Cells were transfected 24 hours post-seeding with pCDNA4/TO, pCDNA4/TO-IE1A, pCDNA4/TO-IE1B expression vector using Lipofectamine 2000 (Thermo Fischer Scientific). After 48 hours of transfection, cells were fixed in 2% paraformaldehyde and used for IFA.

### Infection assays

U2OS cells were seeded at 2 × 10^5^ cells/well in a 6-well plate containing glass coverslips in 2 mL of medium. Cells were infected 24 hours post-seeding with U1102 (HHV-6A) and Z29 (HHV-6B). After 48 hours post-infection, cells were washed 3 times with PBS and fixed in 2% of paraformaldehyde and used for immunofluorescence (IFA) assay.

### Generation of PML Knockout cell line

HeLa LT and U2OS cells were transfected with CRISPR-Cas9 vector targeting PML as described. After 48 hours, cells were selected with 1 µg/mL of puromycin. Selected cells were harvested, counted and seeded at a density of 1 cell per well in three 96-well flat-bottom plates. After 10 to 14 days, wells containing only a single clone were identified. Clones were propagated for an additional 3 weeks and transferred into wells of a 12-well plate. Clones were screened by PCR, sequenced and analyzed by IFA for PML expression. PML negative clones were expanded and kept frozen until used.

### HHV-6A/B integration assays

Integration assays were performed as described previously (45). Briefly, ten thousand cells per well (U2OS PML WT, U2OS PML^-/-^ #1, U2OS PML^-/-^ #2, HeLa LT PML WT, HeLa LT PML^-/-^ #1, HeLa LT PML^-/-^ #2) were seeded in 48-well plates. The next day, the cells were infected with U1102 or Z29 at a multiplicity of infection (MOI) of 1 followed by overnight incubation at 37°C. Cells were washed 3X with 1X PBS to remove unabsorbed virions prior to the addition of fresh culture medium. Upon infection, cells were passaged for 4 weeks and analyzed by droplet digital PCR (ddPCR). For this, DNA was isolated using the QiaAMP blood extraction kit as described by the manufacturer (Qiagen Inc., Toronto, ON, Canada).

### qPCR

qPCR was performed as described previously by Gravel et al. (45). Briefly, DNA was extracted using QiaAMP blood extraction kit as described by the manufacturer (Qiagen Inc.) and analyzed using primers and probes against *U65-66* (HHV-6A/B) and *RPP30* (reference gene). Data was normalized against the corresponding genome copies of the cellular *RPP30* protein.

### Quantification of HHV-6A/B integration by droplet digital PCR (ddPCR)

The HHV-6A/B copy number per cell was determined by ddPCR as previously described by Sedlak et al. (46).

### Statistical analysis

Unpaired t-test with Welch correction was used to compare the number of PML-NBs at telomeres in IE1 expressing and control cells. It was also used to compare the number of IE1 at telomeres in PML^+ / +^ and PML^-/-^ cells. Chi-square analysis was used to compare integration frequency between PML^+ / +^ and PML^-/-^ cell lines.

## Results

### IE1A/B localize at the site of integration, the telomeres

Upon cell entry, HHV-6A/B can either actively replicate or establish latency. This decision is often influenced by the permissivity of the target cells. We have previously shown that U2OS and Hela cells are semi-permissive to infection, as the HHV-6A/B initiates replication in only a minority of cells despite considerable expression of IE and E proteins (45), (47). Both cell lines have been extensively used to assess HHV-6A/B integration (48), (49), (50). To determine if IE proteins might contribute to HHV-6A/B integration, we first determined whether they localize to sites of integration, the telomeres. U2OS cells were infected with HHV-6A (U1102) or HHV-6B (Z29) for 2 days and analyzed for IE1 expression by confocal microscopy. IE1 was detected as distinct nuclear foci upon infection (Fig 1A), with a proportion of IE1 localizing with telomeres (yellow asterisks). Quantification of Z stacks revealed that 20.4% and 26.38% of the IE1A/B foci (red) localize with telomeres (Fig 1B). To assess if IE1A or IEB localize with cellular or viral telomeres, we transfected U2OS cells with IE1A/B expression vectors and analyzed IE1 localization in the absence of viral genomes. Ectopically-expressed IE1A and IE1B localized with cellular telomeres to the same extent as during infection (Fig 2A and B), indicating that cellular telomeres were targeted by these viral proteins.

**Fig 1.**
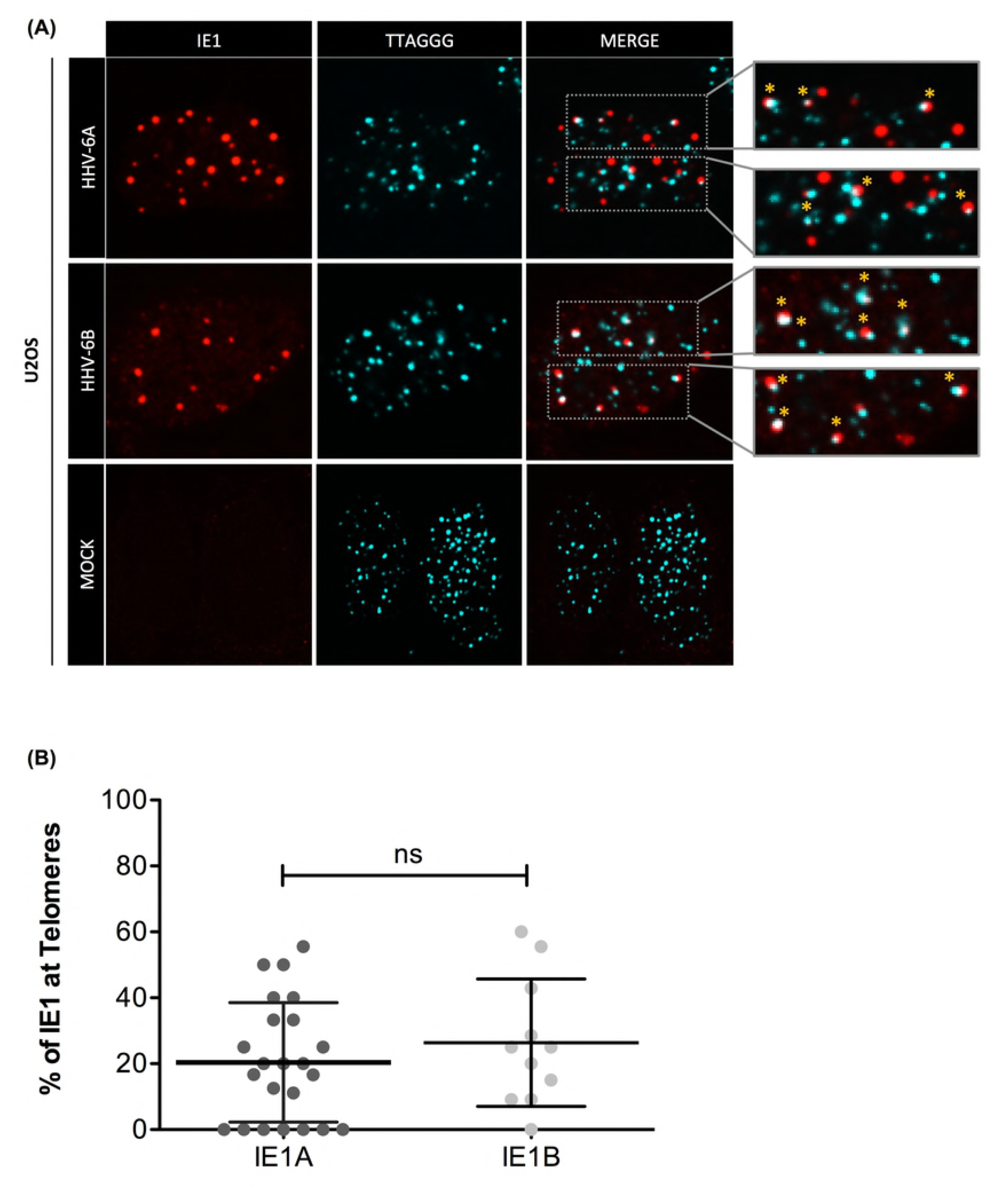
Colocalization of IE1A/B at telomeres in infection. U2OS cells were infected with U1102 (HHV-6A) and Z29 (HHV-6B). A) 48 hours post-infection, cells were fixed with 2% paraformaldehyde and labeled for IF-FISH. IE1A/B was detected using anti-IE1-ALEXA-568 (red) labeled antibodies and telomeres were detected using a Cy5-labeled telomeric probe (Aqua). B) Percentage of IE1A/B foci localizing at telomeres in infected cells. P value was determined using an unpaired t-test with Welch correction. ns: p value is not significant.

**Fig 2.**
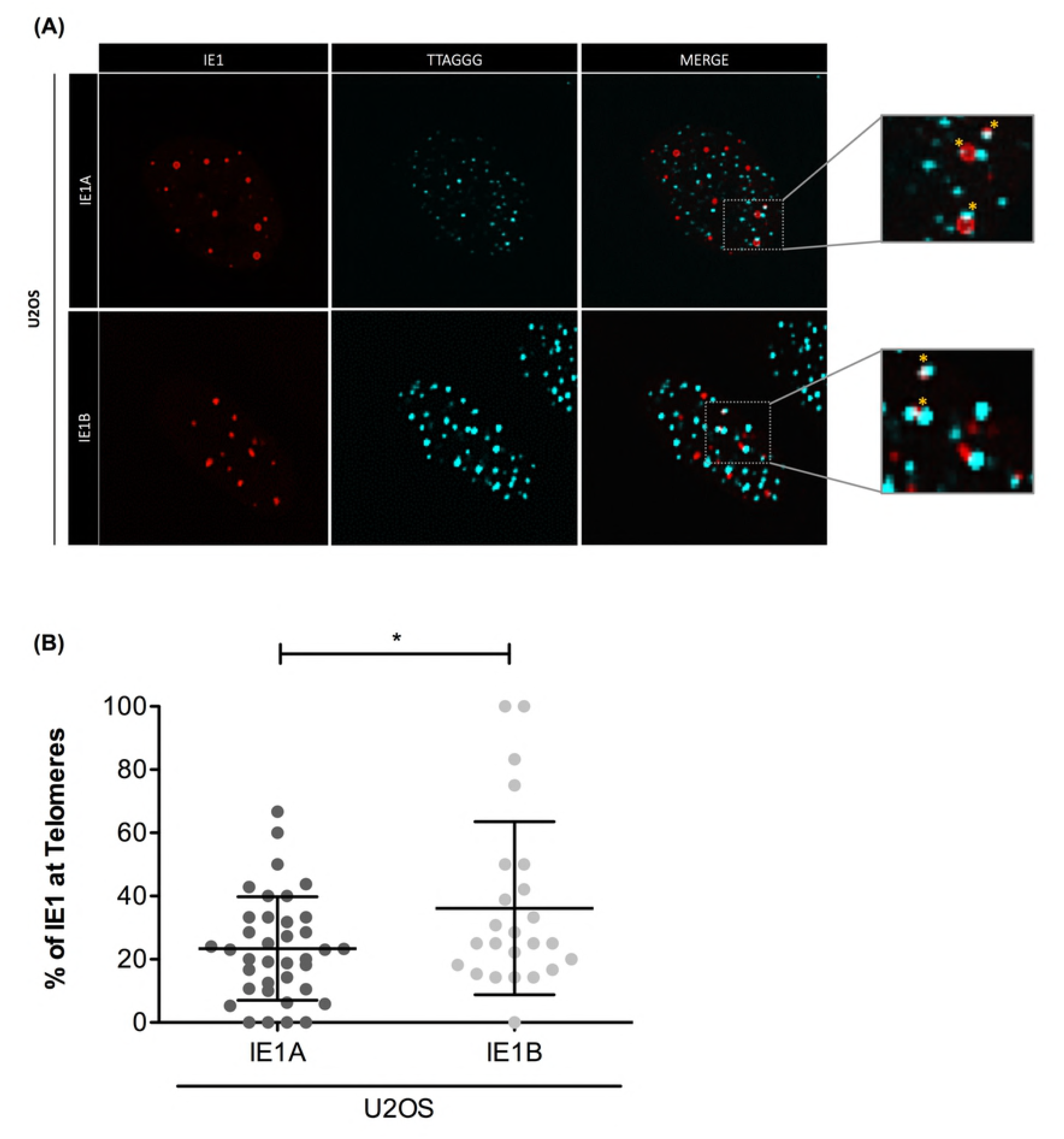
Colocalization of IE1A/B at telomeres in transfection. (A) U2OS cells were transfected with pCDNA4TO (CTRL) and pCDNA4TO-IE1A/B expression vectors. Cells were analyzed by immunofluorescence (IFA) 48 hours post-transfection, using anti-IE1 ALEXA-568-labeled (red) and anti-PML ALEXA-488-labeled antibodies (green). (B) Percentage of IE1A/B foci localizing at telomeres in transfected cells. P value was determined using an unpaired t-test with Welch correction. *P<0.04.

### Both IE1A and IE1B colocalize with PML

We previously demonstrated that IE1 of HHV-6B associates with PML-NBs during productive T cell infection (31). We next determined IE1A and IE1B colocalization with PML-NBs would also be observed in semi-permissive cells. IE1A and IE1B expression vectors were transfected in U2OS cells and their localization was assessed by IFA. IE1 from both viruses efficiently associated with PML (Fig 3).

**Fig 3.**
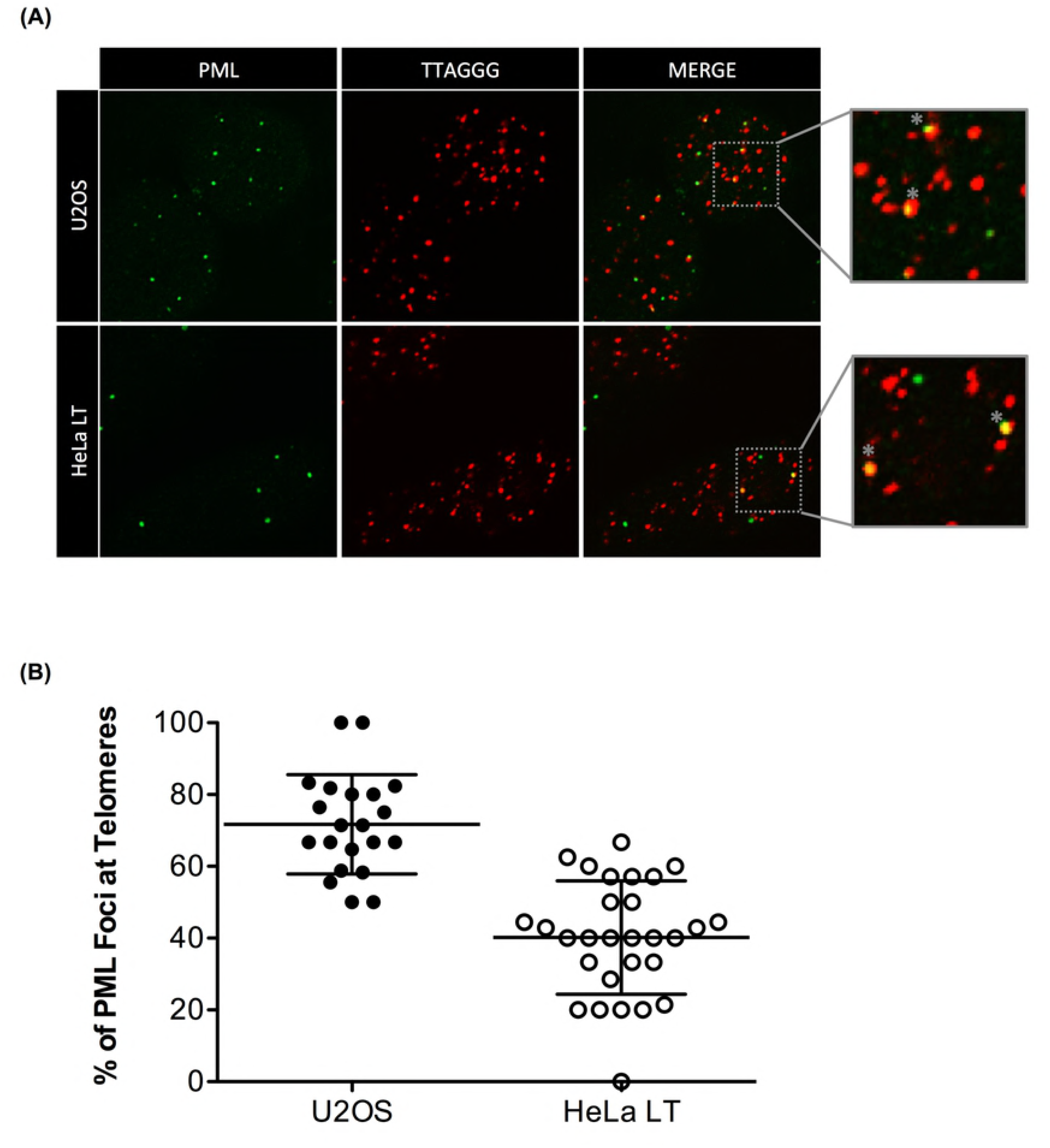
Ectopic IE1A/B colocalize with PML. U2OS were transfected with IE1A/B expression vectors. 48 hours post-transfection, cells were analyzed by IFA using anti-PML ALEXA-488-labeled (green) and anti-IE1 ALEXA-568-labeled (red) antibodies.

PML is actually not a single protein but a mixture of seven different isoforms, whereby the first six isoforms (I to VI) are nuclear proteins (51). To determine if IE1 preferentially colocalizes with certain PML isoforms, we co-transfected PML negative (PML^-/-^) cells with individual expression plasmids for the six nuclear PML isoforms and IE1 expression vectors. Western blotting confirmed that all six PML isoforms are efficiently expressed upon transfection of PML^-/-^ cells (Fig 4A). IE1B colocalized with all PML isoform tested. Similarly, IE1A localized with all 6 PML isoforms (data not shown).

**Fig 4.**
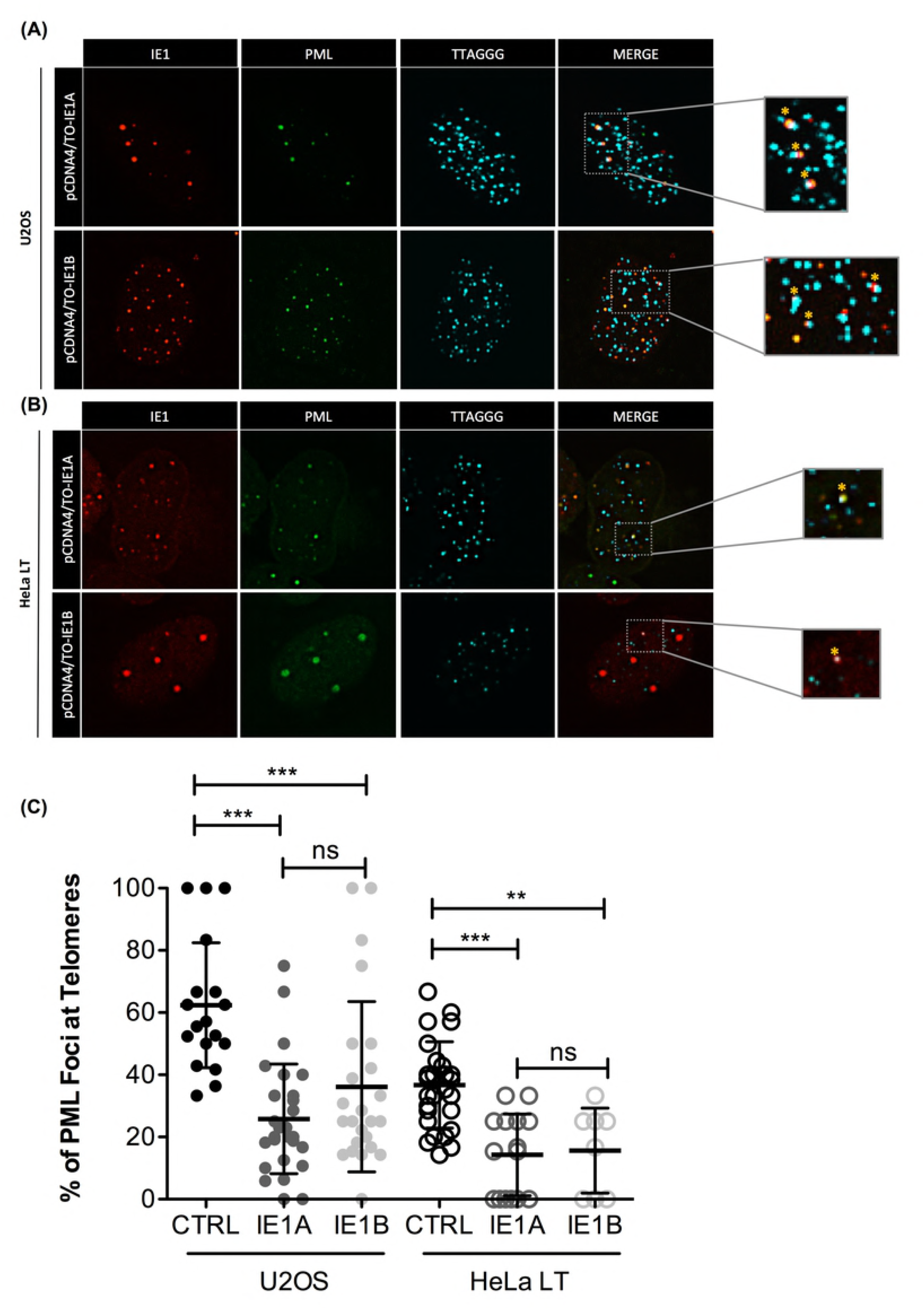
IE1A/B colocalize with all nuclear PML isoforms. (A) HEK293T were transfected with PML isoforms expression vectors and analyzed by western blot using anti-Myc antibodies. U2OS PML^-/-^ were co-transfected using pCDNA4TO-IE1B (B) vectors along with vectors expressing the various PML isoforms (I to VI). 48 hours post-transfection, cells were analyzed by IFA using anti-Myc ALEXA-488-labeled (green) and anti-IE1 ALEXA-568-labeled (red) antibodies.

### Presence of IE1A/B affects the number of PML-NBs present at cellular telomeres

U2OS cells do not express telomerase and elongate their telomeres via alternative lengthening of telomere mechanisms (ALT) (52), (41), (53). In ALT cells, a significant proportion (75%) of PML-NBs localize at telomeres and are referred to as ALT-associated PML-NBs (APBs) (Fig 5) (54). In Hela LT cells that rely mostly on the telomerase complex for telomere elongation the number of APBs was much lower (Fig 5).

Since IE1A/B colocalize with PML-NBs, we next assessed whether IE1’s presence might affect PML-NBs localization at telomeres. U2OS and HeLa LT cells were transfected with IE1A/B expression vectors, and the proportion of PML-NBs localizing to telomeres was determined by IF-FISH (Fig 6A and B). The frequency of PML-NBs located at the host telomeres was reduced by 58% (25.8+/- 22.53) in U2OS cells upon expression of IE1A compared to the empty vector control (63.35+/- 16.97) (Fig 6C). A comparable reduction in APBs of 50% was also observed in U2OS cells expressing IE1B. We confirmed this observation in HeLa LT cells, where the PML-NBs localizing at telomeres was reduced by 64% and 61% upon expression ofIE1A and IE1B respectively. Similar results were obtained in U2OS and HeLa LT cells infected with HHV-6A/B (data not shown).

**Fig 5.**
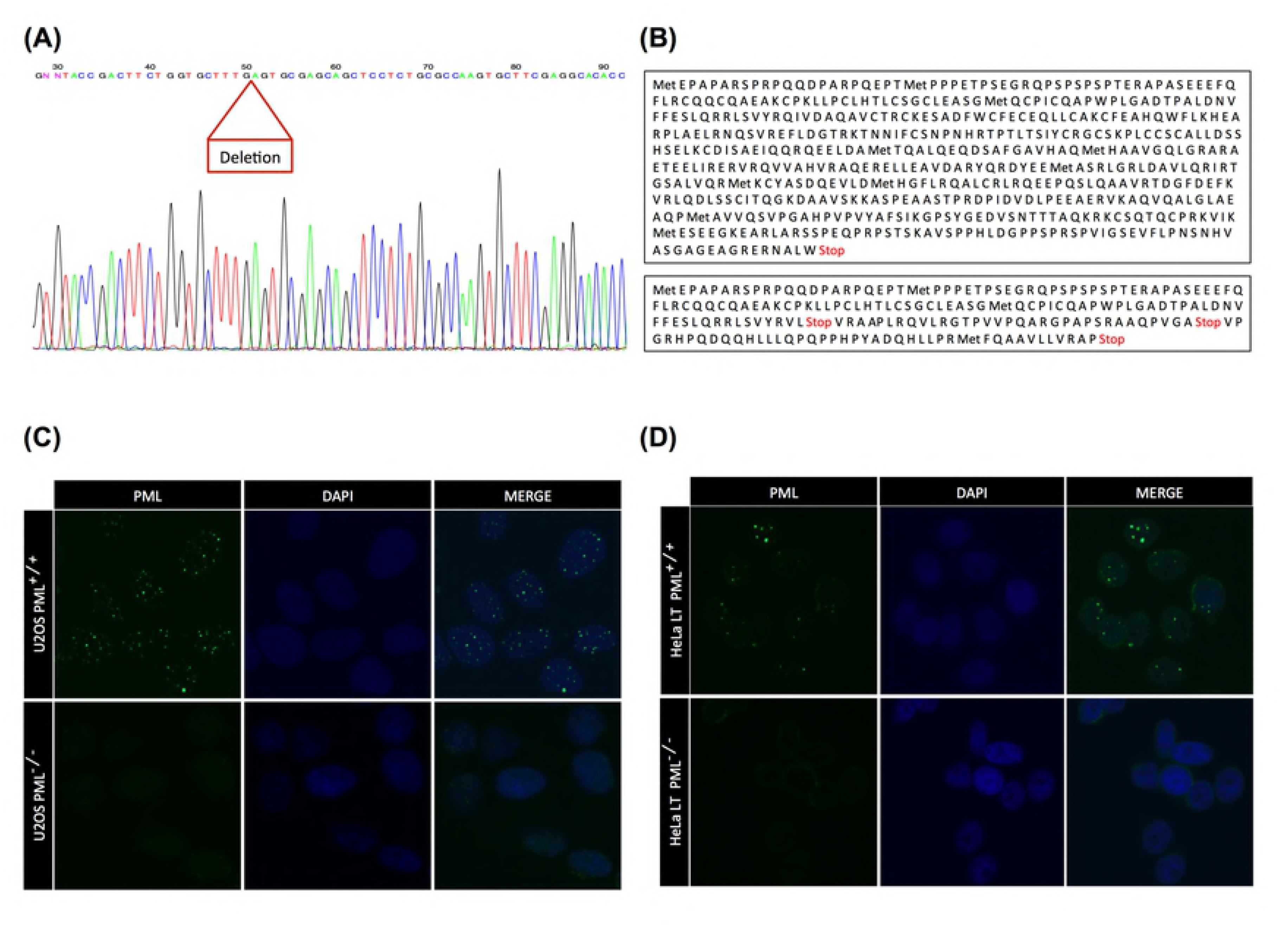
PML proteins colocalize at telomeres in U2OS and HeLa LT cells. (A) U2OS cells (ALT+) and HeLa LT cells (telomerase +) were grown on coverslips and fixed with 2% paraformaldehyde at sub confluence. Cells were analyzed by IF-FISH. PML proteins were detected using anti-PML ALEXA-488-labeled (green) antibodies and telomeres were detected using a Cy3-labeled telomeric probe (red). (B) The number of PML foci localizing at telomeres was calculated after analysis of U2OS (N=20) and HeLa LT (N=40) nuclei.

**Fig 6.**
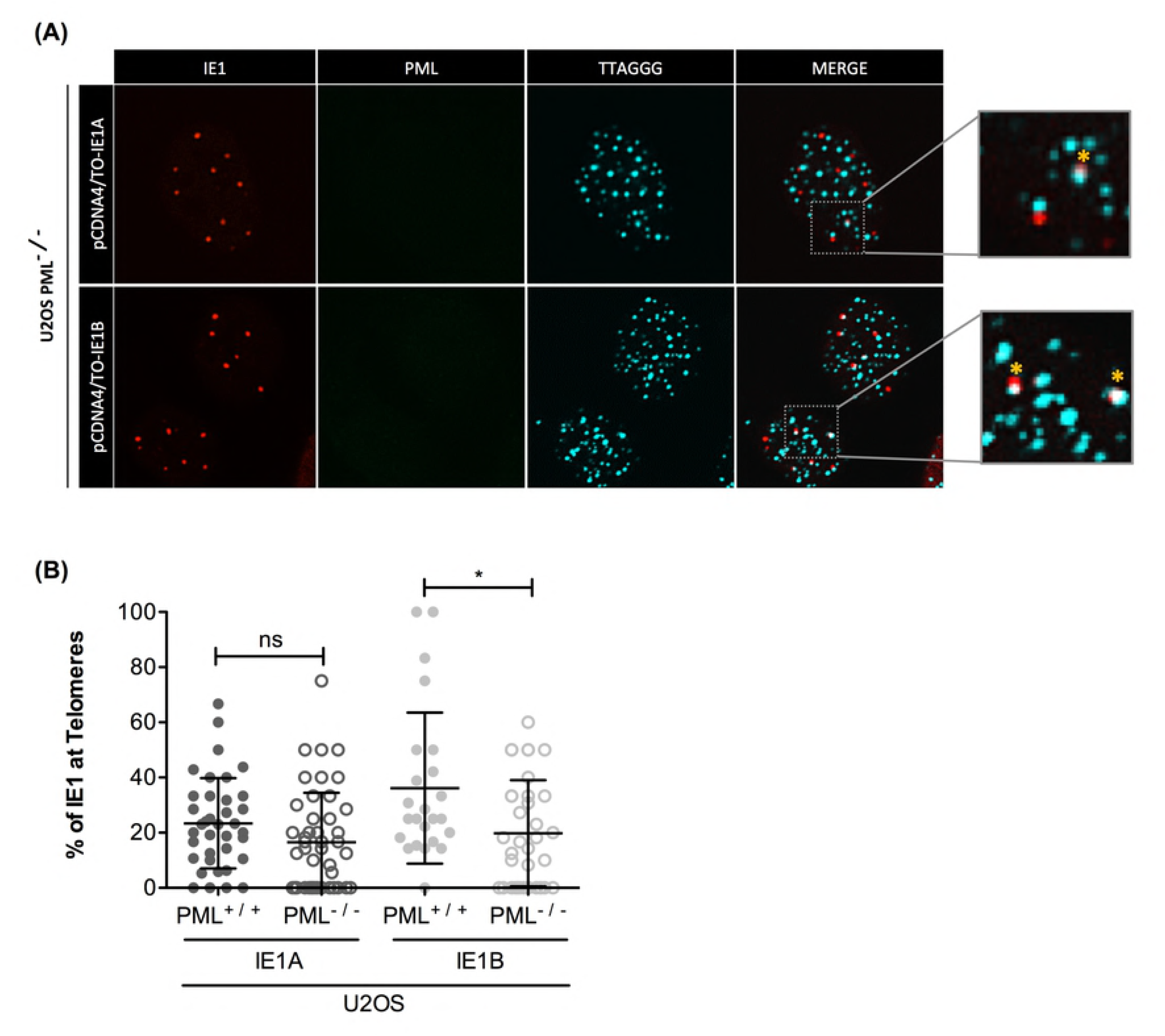
IE1A/B colocalize with PML at telomeres. U2OS (A) and HeLa LT (B) cells were transfected with pCDNA4/TO-IE1A or pCDNA4/TO-IE1B. 48 hours post-transfection, cells were fixed with 2% paraformaldehyde and analyzed by IF-FISH. IE1A/B were detected using anti-IE1-ALEXA-568-labeled antibodies (red), PML were detected using anti-PML ALEXA-488-labeled (green) antibodies and telomeres were labeled using a Cy5-labeled telomeric probe (Aqua). (C) Percentage of IE1A/B at telomeres in transfected U2OS (N=37) and HeLa LT (N=24) cells. P value was determined using an unpaired t-test with Welch correction. *P<0.01; ns = p value is not significant. (D) Percentage of PML foci at telomeres in presence and in absence of ectopically expressed IE1A/B. CTRL: Empty vector. P value was determined using an unpaired t-test with Welch correction. ***p<0.0001

### The absence of PML does not affect the presence of IE1A/B at telomeres

Considering that IE1A/B colocalize with PML-NBs and that a significant proportion of PML-NBs are located at host telomeres, we next determined if PML contributes to the localization of IE1A/B’s to the telomeres. PML knockout (KO) U2OS and HeLa LT cell were generated using the CRISPR-Cas9 system (Fig 7). Deletion of a part of exon 1 (Fig 7A) resulted in a pre-mature STOP codon resulting in a short truncated PML protein (Fig 7B). Abrogation of PML expression was confirmed in U2OS (Fig 7C) and HeLa LT (Fig 7D) cells by IFA.

Following transfection of IE1A/B expression vectors in WT and PML^-/-^ U2OS cells, we observed that IE1A/B localized at telomeres despite PML’s absence, albeit at a slightly lower frequency (Fig 8A and B). There was however an increased proportion of cells in which IE1A/B were not present at telomeres. As shown in Fig 8C, the number of U2OS PML^-/-^ nuclei with no IE1A/B at telomeres was significantly increased relative to WT nuclei (***p<0.0001).

**Fig 7.**
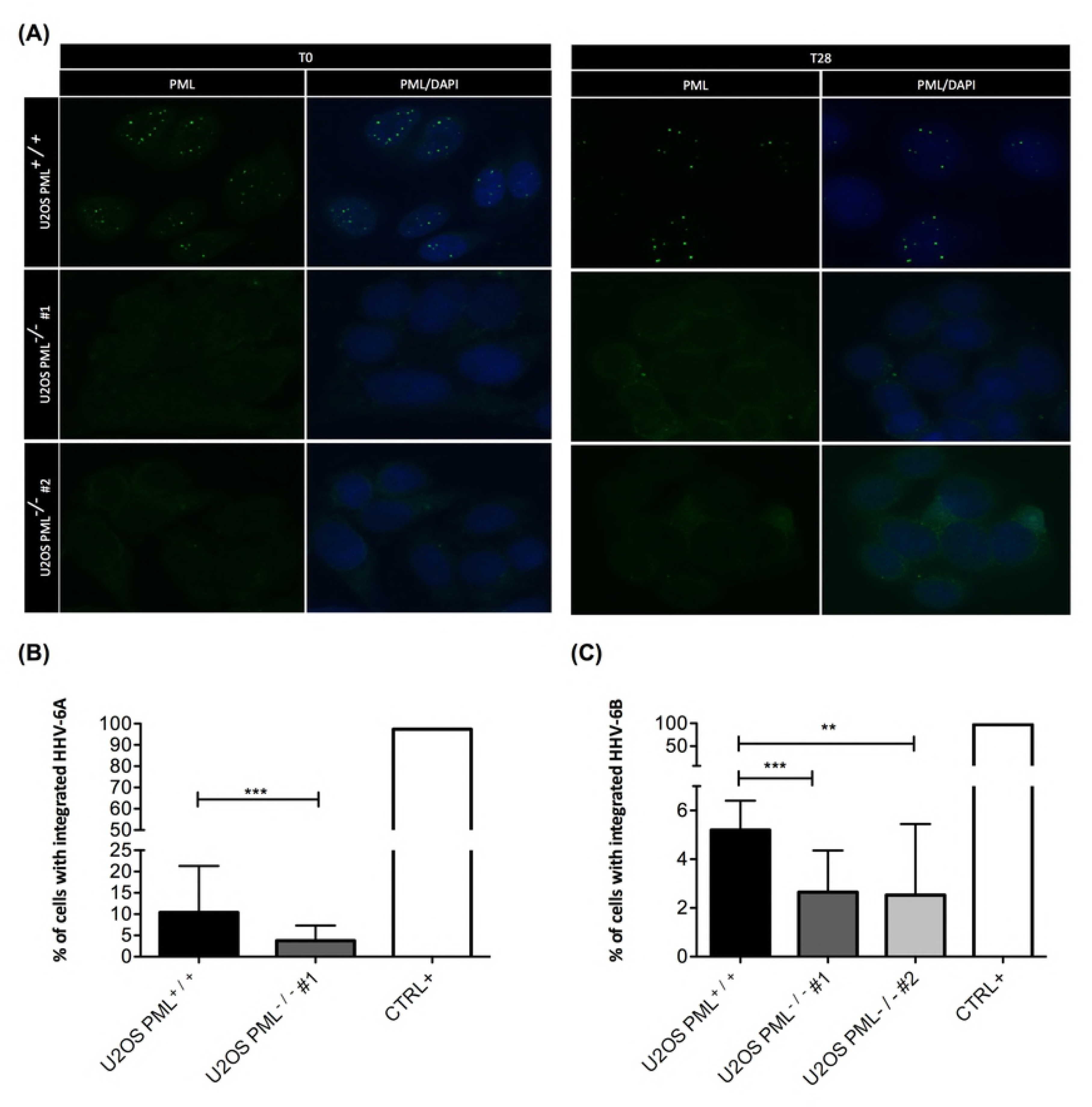
Generation of PML Knockout cell lines. U2OS and HeLa LT cells were transfected with expression vectors for Cas-9 nuclease and PML RNA guides. After puromycin selection, cells were seeded at a density of one cell/well to obtain unique clones. (A) Each clone was screened by PCR with PML primers. Mutations were confirmed by sequencing the PCR amplicons. (B) Translation of the mutated sequence results into a truncated protein with three premature STOP codons. WT and PML^-/-^ U2OS (C) and HeLa LT (D) cells were analyzed by IFA for PML expression using anti-PML ALEXA-488-labeled (green) antibodies.

**Fig 8.**
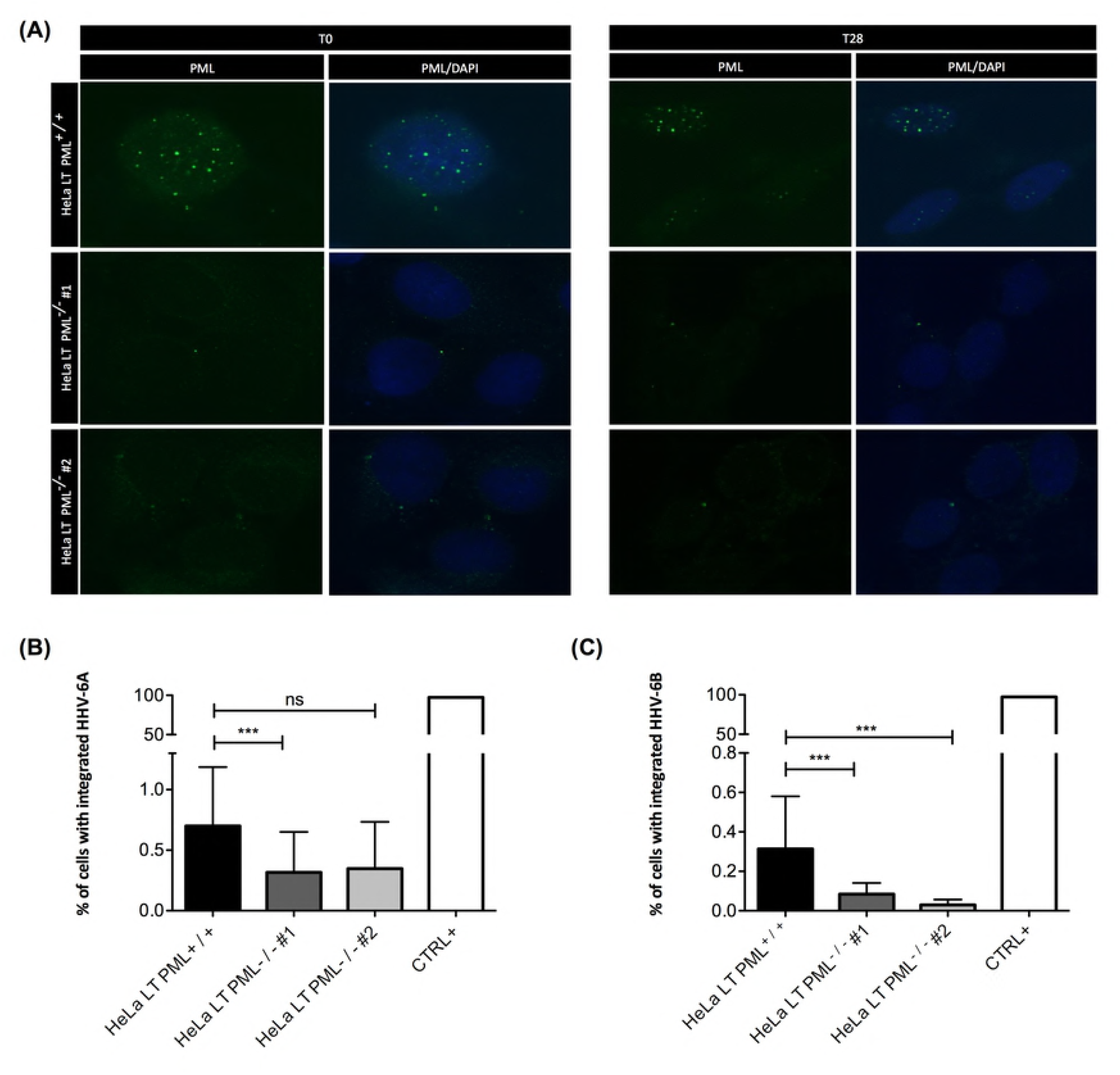
PML is dispensable for IE1A/B localization at telomeres. (A) U2OS^-/-^ cells were transfected with pCDNA4TO-IE1A and pCDNA4TO-IE1B expression vectors. 48 hours post-transfection, cells were analyzed by IF-FISH for IE1A/B (red) and PML (green) expression using specific antibodies. Telomeres were detected using Cy5-labeled telomeric probe (Aqua). (B) Number of IE1A/B foci localizing at telomeres in the presence (N=37 for IE1A PML^+ / +^; N=24 for IE1B PML^+ / +^) or in the absence of PML (N=46 for IE1A PML^-/-^; N=35 for IE1B PML^-/-^). P value was determined using an unpaired t-test with Welch correction. *P<0.01. ns: p value is not significant. (C) Total number of IE1A/B that have no IE1A/B at telomeres was compared between PML^+ / +^ (N=37 for IE1A; N=24 for IE1B) and ^-/-^ cell lines (N=46 for IE1A; N=35 for IE1B). P value was determined using Chi-square analysis. ***P<0.0001

### PML is required for efficient HHV-6A/B chromosomal integration

Considering that a proportion of PML-NBs localize at telomeres and that PML plays a role in DNA repair by homologous recombination, we investigated if PML plays a role in HHV-6A/B integration into telomeres. PML knockout and control cells lines were infected with HHV-6A or HHV-6B and integration frequency assessed after four weeks post infection by droplet digital PCR as described (48). The absence of PML was confirmed at the beginning (T0) and the end (T28) of the experiment by IFA for U2OS (Fig 9A) and Hela LT cells (Fig 10A). ddPCR revealed that HHV-6A and -6B integration was significantly reduced in U2OS cells in the absence of PML by 64% and 50% respectively (Fig 9B and C). In Hela LT cells, HHV-6A integration was reduced by approximately 50% in the absence of PML (Fig 10B). The reductions were even more pronounced for HHV-6B in HeLa LT cells where the integration frequency was reduced by 73% and 90.6% in the two independent clones used in this study (Fig 10C). Taken together, our data demonstrates that integration occurs less efficient in the absence of PML in two standard models for HHV-6A/B integration.

**Fig 9.**
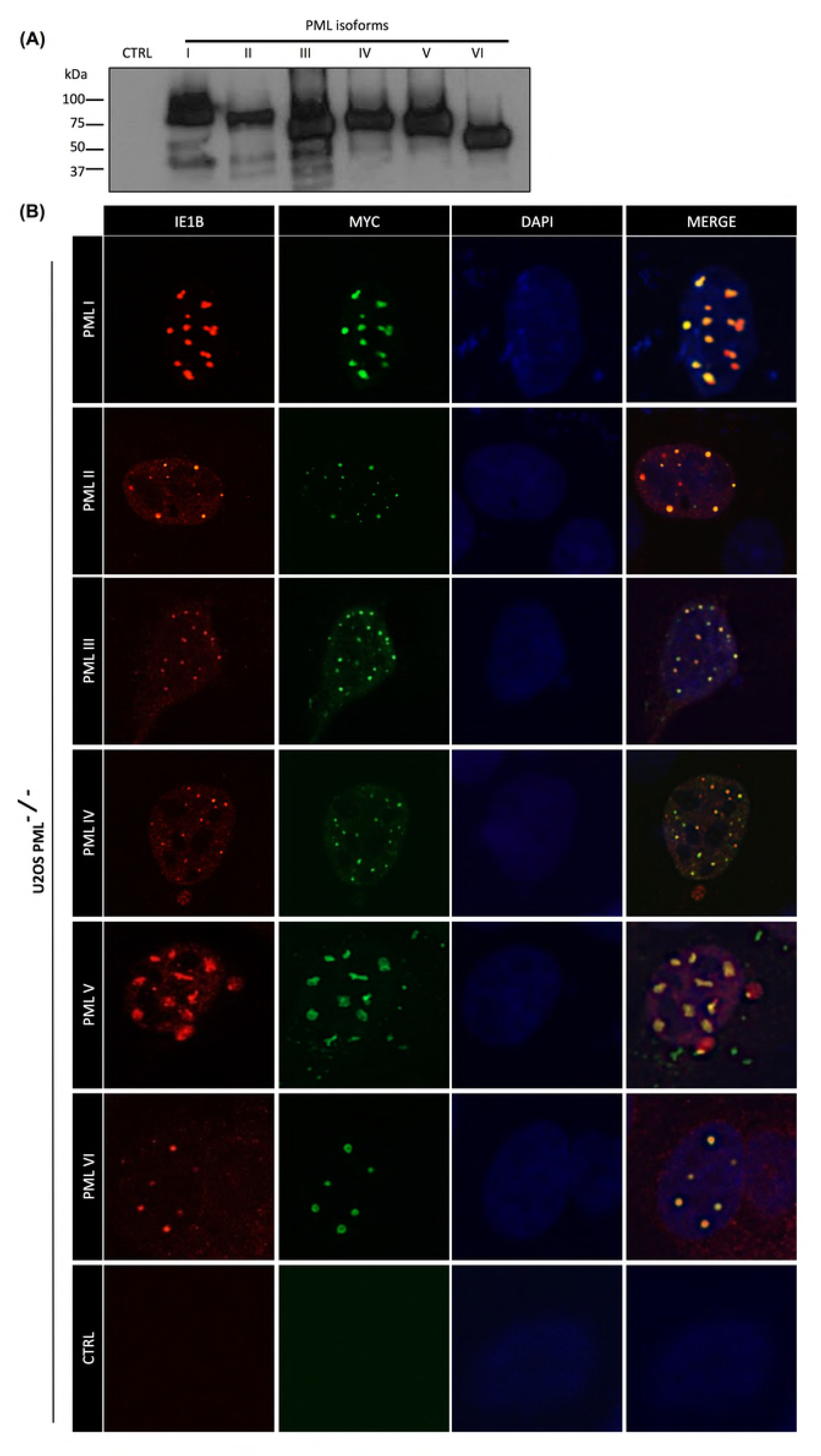
HHV-6A/B integration in WT and PML ^-/-^ U2OS cells. (A) PML expression in WT and PML^-/-^ U2OS cells on day 0 and day 28 post-infection. WT and PML^-/-^ U2OS cells were infected at a MOI of 1 with HHV-6A (B) and HHV-6B (C) and were cultured for a month. Cellular DNA was extracted, and integration frequency determined by ddPCR. Each integration assay was done three time for each cell lines (error bars). CTRL +: iciHHV-6A/B donor DNA. P value was determined using Chi-square analysis. P value was determined using Chi-square analysis. ***P<0.0001; **P<0.001

**Fig 10.**
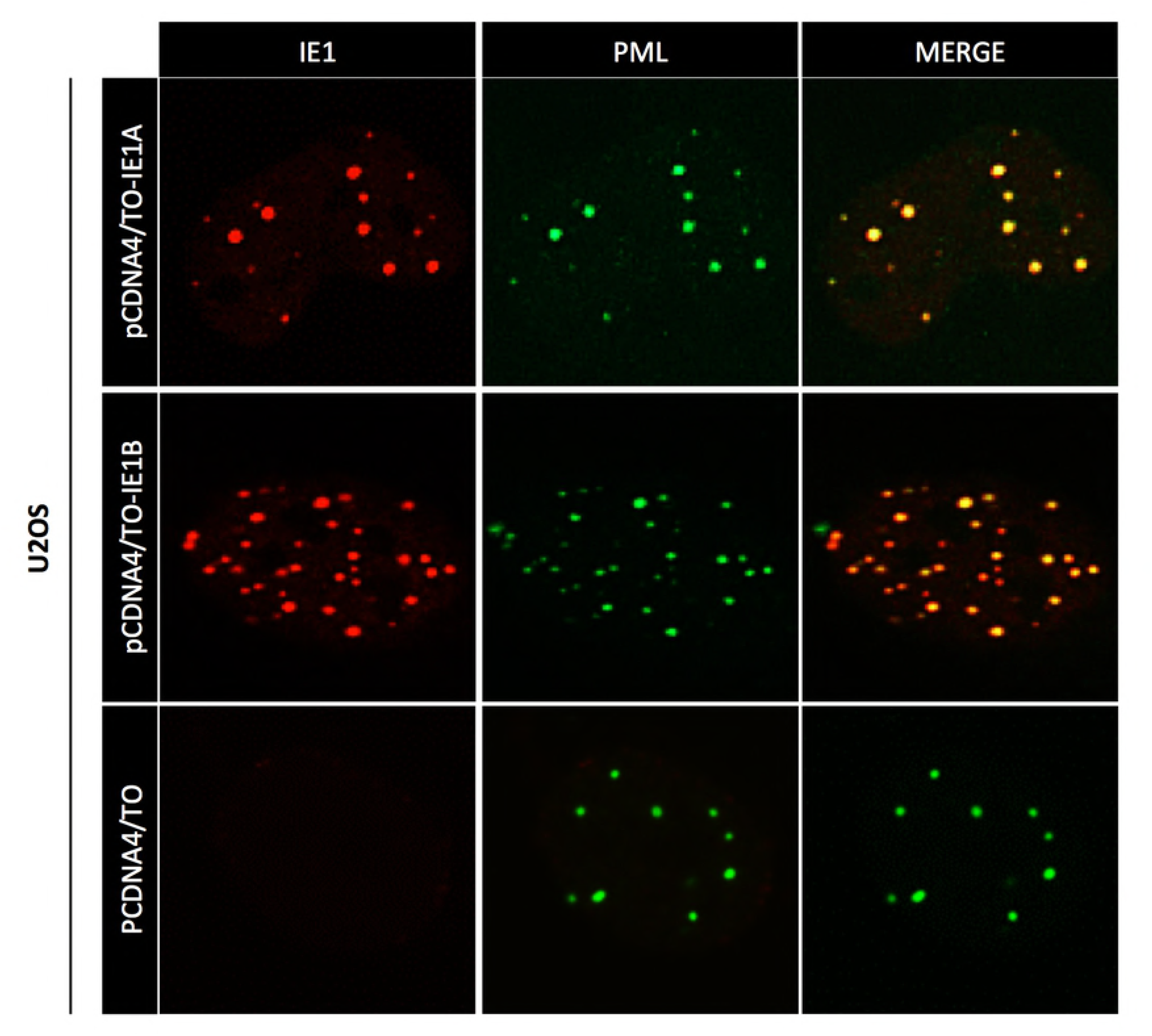
HHV-6A/B integration in WT and PML ^-/-^ HeLa LT cells. (A) PML expression in WT and PML^-/-^ HeLa LT cells on day 0 and day 28 post-infection. WT and PML^-/-^ HeLa LT cells were infected at a MOI of 1 with HHV-6A (B) and HHV-6B (C) and were cultured for a month. Cellular DNA was extracted, and integration frequency determined by ddPCR. Each integration assay was done three time for each cell lines (error bars). CTRL +: iciHHV-6A/B donor DNA. P value was determined using Chi-square analysis. ***P<0.0001; ns: p value is not significant.

## Discussion

One key interest of our laboratory is to identify proteins that facilitate HHV-6A/B chromosomal integration. We previously demonstrated that the putative HHV-6A/B integrase U94 possesses DNA binding, helicase and exonuclease activity, suggesting that the protein could be involved in HHV-6A/B integration (55). However, recombinant HHV-6A lacking U94 integrated as efficient as WT virus, indicating that U94 is dispensable for integration of HHV-6A in vitro (50).

Another hypothesis is that HHV-6A/B chromosomal integration occurs with the help of telomerase, the enzyme responsible for telomere elongation (56). We have previously shown that telomerase is not essential for HHV-6A/B integration, as it occurs in both telomerase negative and positive cells (48), (49). However, in telomerase expressing cells, telomerase is likely important for the generation a neo-telomere at the end of DR_L_ (reviewed in (57)). In support, blockade of telomerase activity by the G-quadruplex (guanine rich structure (G4) present in telomeres) stabilizing agent BRACO-19, negatively affects HHV-6A integration (49). Such effect is not observed in telomerase negative cells such as U2OS cells.

The fact that HHV-6A telomeric repeats are required for efficient integration (30) argues in favor of a homologous recombination (HR) events between cellular and viral telomeres. Cellular telomeres are protected by the shelterin complex that consists of 6 proteins: telomeric repeat binding factor 1 (TRF1), TRF2, protection of telomere 1 (POT1), telomere protection protein 1 (TPP1), TRF-interacting nuclear protein 2 (TIN2) and repressor activation protein 1 (RAP1) (58). The main function of the shelterin complex is to protect chromosome end from being recognized as damaged DNA. Of the 6 shelterin proteins, TRF2 is the key factor that blocks DNA repair proteins at telomeres by inhibiting the Ataxia-telangiectasia-mutated (ATM) pathway that senses double-stranded DNA breaks (59). In addition to the shelterin complex, other proteins can also localize to telomeres. In telomerase negative cells such as U2OS cells, telomeres are elongated by an Alternative Lengthening of Telomeres (ALT+) associated PML-Nuclear Bodies (APBs) mechanism (60),(54),(61),(62). These nuclear bodies primarily formed by the Promyelocytic Leukemia Protein (PML) itself that recruits hundreds of interacting partners at telomeres such as helicases implicated in G-quadruplex structure resolution like the bloom syndrome protein (BLM), the Werner Syndrome Protein (WRN) and other protein implicated in DNA recombination and repair (63), (54), (64), (65), (53). Osterwalds et al. have shown that in ALT^+^ cells such as U2OS, PML-NBs (APBs) are frequently present at telomeres. We’ve confirmed this result (Fig 5). We also noticed that a significant proportion of PML-NBs also localize to telomeres of telomerase expressing cells such as HeLa LT cell (Fig 5), in agreement with Marchesini et al (63). Marchesini demonstrated that PML is essential for telomere maintenance in non-neoplastic cells, as cells undergo apoptosis in absence of PML after DNA damage at these sites (63). These findings support the role of PML in DNA repair mechanism. Considering this, we hypothesized that PML could aid in HHV-6A/B integration.

In addition to their roles in telomere stability, PML-NBs have antiviral defense functions (36), (35). In contrast to many other viruses including HSV, CMV, EBV and HHV-8, HHV-6A/B infection does not lead to the dispersal of PML-NBs but rather to PML-NBs coalescence (38), (66), (67), (68), (33). Whether this affects antiviral functions of PML-NBs remains to be determined. We could demonstrate that the HHV-6A/B IE1 protein, a protein that play roles in innate immune evasion mechanisms (69), (42), is associated with PML-NBs (31). Here we also report that IE1A/B also associates with telomeres. Considering that PML also associates with telomeres, we hypothesized that IE1A/B localization to telomeres could be PML dependent. However, in PML^-/-^ U2OS and HeLa LT cells, a significant proportion of IE1A/B remained associated with telomeres. However, the proportion of nuclei in which IE1A/B could not be detected at telomeres was largely increased in PML^-/-^ cells. Thus, although not essential, PML does influence the localization of IE1A/B at telomeres. One possible explanation resides in the fact that IE1A/B are SUMOylated proteins and that PML itself and/or other PML-NB associated proteins contain SUMO interacting motif (SIM) could facilitate interactions at telomeres (31), (70). Moreover, IE1A/B also possess putative SIMs that can bind SUMOylated proteins present at telomeres, possibly explaining why IE1A/B can localize at telomeres in the absence of PML.

Finally, we tested if PML played a role in HHV-6A/B chromosomal integration. We used the CRISPR-Cas9 to abrogate PML expression (Fig 7). For each cell line used, two independent PML KO clones were assessed to ensure reproducibility and avoid potential off-target effects. In U2OS cells (Fig 9), HHV-6B integration was less frequent in PML^-/-^ cells (p<0.0001). In HeLa LT cells (Fig 10), the same effect was observed for HHV-6A and HHV-6B. Of note, integration rates are higher in U2OS presumably because of higher constitutive DDR repair in the cells, supporting the hypothesis that DNA repair mechanisms are involved in the HHV-6A/B integration process (71). Globally, both cell lines studied suggest a role for PML in HHV-6A/B integration. However, since integration still occurred in PML KO cells indicates that PML contributes but is not essential for this process. The positive influence of PML on HHV-6A/B integration could be explained by a reduction of protein present at telomeres like those involve in the DDR. As mentioned above, TRF2 blocks DNA repair at double DNA strand breaks. Moreover, in PML-NBs at telomeres, TRF2 is SUMOylated by MMS21, resulting in a lower density of TRF2 on telomeres (72). These telomere regions can then be processed by other proteins and recombine with the HHV-6A/B telomeres.

In conclusion, we have demonstrated that HHV-6A/B IE1 proteins colocalize with all isoforms of PML and host telomeres. Abrogation of PML expression influences the presence of IE1 at telomeres and affects HHV-6A/B integration into host telomeres. To our knowledge, this is the first report of a cellular protein that is involved in HHV-6A/B integration.

## Supporting information

**S1 Fig. PML KO does not create more DNA damages.** PML^+ / +^ and PML^-/-^ cells from the integration assays at T28 were seeded on coverslips and fixed with 2% of paraformaldehyde. Cells were analyzed by IF-FISH for DNA damage protein 53BP1 (red) and PML (green) expression using specific antibodies. Telomeres and nuclei were detected using Cy5-labeled telomeric probe (Aqua). Number of 53BP1 foci per nuclei was counted for PML^+ / +^ U2OS cells (N=42), HeLa LT cells (N=37) and PML^-/-^ U2OS cells (N=40) and HeLa LT cells (N=37). P value was determined using an unpaired t-test with Welch correction. ns: p value is not significant.

**S2. PML and IE1B localize in close proximity.** U2OS-Flp-In TREX cells were transfected with expression vectors containing FLAG-BirA-GFP and FLAG-BirA-IE1B and selected with hygromycin (250µg/ml) and blasticidin (50µg/ml). (B) Cells were seeded on coverslips and 24 hours later, 50nM of biotin was added to the medium for an additional 24h before being fixed with paraformaldehyde 2%. IFA confirms BirA-GFP and BirA-IE1B expression (Flag) and biotinylation of proteins (Streptavidin-HRP). (C) Biotinylated proteins were immunoprecipitated with streptavidin magnetic beads followed by mass spectrometry.

